# Survivorship Bias Explains Age-Dependent Extinction in Fossil Genera

**DOI:** 10.64898/2026.07.21.739780

**Authors:** Mikhail Tikhonov, Kim Sneppen, Stefan Bornholdt, Sergei Maslov

## Abstract

Fossil genera exhibit pronounced age-dependent extinction: older genera are more likely to survive extinction events than younger ones. Many biological mechanisms have been proposed that could make lineages progressively more resistant to extinction through time. However, the same pattern can also arise from survivorship bias: as more vulnerable genera are progressively eliminated, the surviving pool becomes increasingly enriched in robust lineages, even if the properties of individual lineages never change. Using the Sepkoski marine genus compendium, we construct a quantitative model based solely on survivorship bias and ask whether additional age-dependent changes are required to explain the fossil record. This model with a single fitting parameter quantitatively reproduces the full set of age-conditioned survival probabilities, together with the overall dependence of extinction risk on genus age and the genus lifetime distribution. Allowing extinction resistance to change systematically through time yields no measurable improvement, indicating that, at least at the aggregate statistical level, explicit age-dependent changes in lineage properties are not required to explain the observed patterns of extinction. These results show that our minimal one-parameter model provides a quantitative null model for age-dependent extinction, against which proposed biological mechanisms can now be tested.

---

Macroevolutionary analyses of the fossil record have long noted that older taxa often appear less likely to go extinct than younger ones [1–9]. Although some planktonic groups show the opposite pattern, with extinction risk increasing with age [10], a broad range of marine clades exhibit elevated extinction among young taxa and suppressed extinction among older ones [9, 11, 12]. The pattern persists even after controlling for geographic range and genus richness, indicating that it cannot be reduced to simple correlates of ecological success [9]. These observations stand in contrast to the expectations of the “law of constant extinction” and the Red Queen hypothesis, which predict that extinction risk should remain approximately independent of lineage age [13].

The explanations proposed for this pattern can be grouped into two broad classes. One invokes biological mechanisms leading to explicit age-dependent changes in extinction resistance. In this view, lineages become harder to eliminate as they persist through time. This can occur through adaptive mechanisms, such as evolutionary refinement or increasing ecological entrenchment, or through neutral processes that gradually increase a taxon’s abundance, occupancy, or diversity [12]. In all such scenarios, older taxa survive better because something about the lineage itself systematically changes with age.

An alternative explanation is survivorship bias arising from heterogeneity in intrinsic robustness [14, 15]. In this view, taxa differ in their ability to withstand environmental and biotic stresses. As the more fragile lineages are progressively removed, the surviving pool becomes enriched in taxa with greater resistance to future perturbations. This mechanism will produce apparent age dependence even if the intrinsic perturbation resistance of each lineage remains constant through time.

It seems inevitable that survivorship bias is responsible for at least part of the observed age dependence in extinction risk. Interpreting the extent to which the fossil record reflects additional biological processes therefore requires understanding how much of the pattern can already be explained by this statistical effect alone. Here, we use the Sepkoski marine genus dataset [7] to show that a simple one-parameter model based solely on survivorship bias reproduces the major statistical features of the fossil record surprisingly well. Furthermore, introducing explicit age-dependent changes in robustness does not improve the agreement with the data. Our results establish a quantitative null expectation against which additional biological mechanisms can be evaluated.

## RESULTS

### A. Survivorship bias is a real effect

The most direct test for survivorship bias is to compare survival probabilities (in a given *focal event*) between matched cohorts of genera whose origination times are separated by a major earlier extinction (*filtering event*; Fig. 1A). The survivorship bias mechanism predicts that such cohorts would exhibit a larger difference in survival probability than expected from age difference alone.

**FIG. 1.**
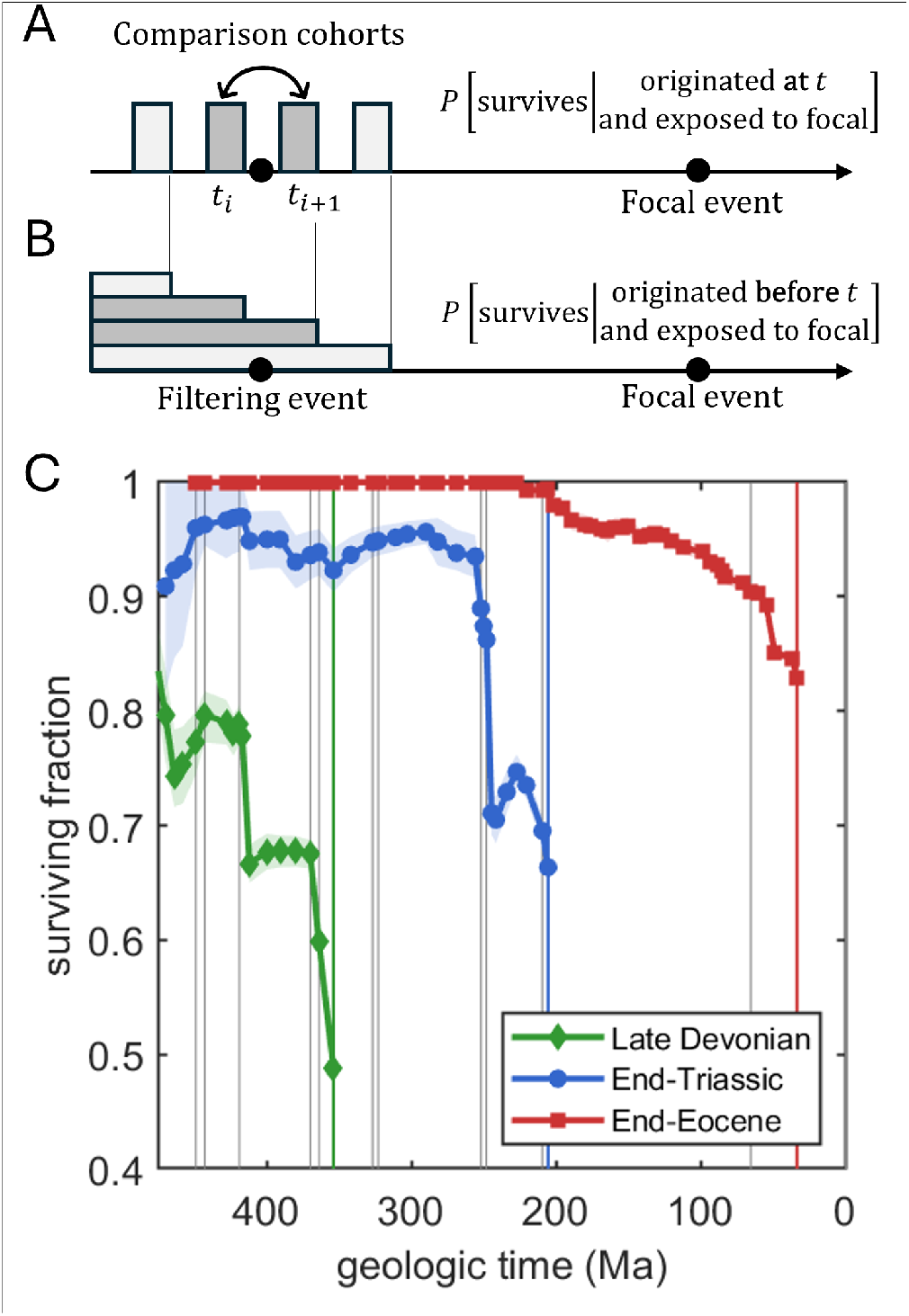
Event survival curves reveal age dependence punctuated by earlier extinction filters. A–B: Rather than defining cohorts as genera originating immediately before or after an earlier filtering event (A), we define them as all genera alive at an earlier time *t* and surviving until the focal event (B; see text). This substantially increases sample size. C: Examples of the resulting event survival curves. Each curve shows, for one focal extinction event, the fraction of the corresponding retrospective cohort that survives that event as a function of *t*. Thus, moving leftward to older times progressively restricts the analysis to older retrospective cohorts. The three examples shown correspond to the Late Devonian, End-Triassic, and End-Eocene events, respectively; the corresponding vertical colored lines mark the focal events. Vertical gray lines mark events with extinction fractions exceeding 40%. Shading indicates the interquartile range (IQR) estimated from 1000 bootstrap resamplings of genera within each cohort; cohorts with < 10 genera are omitted. The curves generally rise for older cohorts, but the sharp jumps near earlier extinction events show that the age dependence is not only a smooth function of age, but bears the imprint of prior extinction filters.

However, this direct approach has limitations. Conceptually, this test is sharpest if the two cohorts are defined as originating just before and just after a filtering event, but the number of old genera surviving until a focal event is often small, and further conditioning on a narrow origination time window makes comparisons very noisy (see supplementary figure S1).

To deal with this statistical issue, we will structure our comparison differently. Our main object will be the *event survival curve*, defined for each event *E* as the conditional probability

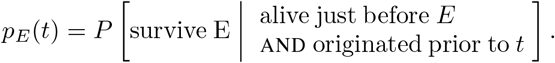

The key is that rather than conditioning on genera that originated within a given stratigraphic epoch (Fig. 1A), we include all genera alive at that time (Fig. 1B). This substantially increases the number of genera contributing to the estimate. The term “event survival curve” emphasizes that this retrospectively-defined quantity is centered on a specific extinction event and describes how the probability of surviving that event depends on lineage history (Finnegan et al. presented an example of such a curve in Fig. 1 of [9]). This contrasts with Raup’s survivorship curves, which are cohort-centered and ask how the descendants of taxa present at a given time persist into the future.

Fig. 1C shows three examples of event survival curves. As expected, survival probability generally increases as we move further into the past, indicating that older genera are more likely to survive the focal event. Crucially, superimposed on this overall trend are pronounced jumps, many of which appear to coincide with earlier major extinction events (events with extinction fractions exceeding 40% are marked by vertical gray lines).

Some discontinuities are expected even under a purely age-dependent extinction process. Major extinction events alter the composition of the surviving cohort and can therefore produce abrupt changes in its average age. However, the corresponding diagnostic curves of mean cohort age (Supplementary Fig. S2) are substantially smoother, which suggests that the jumps in the event survival curves are not solely a consequence of changes in age structure. This observation is consistent with survivorship bias. Compared to the narrow-window analysis of Fig. 1A and Fig. S1, the larger statistics allow event survival curves to be estimated with lower uncertainty. This makes event survival curves an attractive object for quantitative model comparison, and they will be our primary focus below. Although the illustrative Fig. 1C shows only three examples, event survival curves can be defined for any epoch boundary. Below, we evaluate the ability of different models to explain the ensemble of 84 such curves constructed from the Sepkoski dataset.

### B. A zero-parameter model with survivorship bias as its only ingredient

To ask how much of the survival-curve structure can be explained by survivorship bias alone, we construct a simple generative model that we will call the toughness threshold model. For each geologic interval *i*, the Sepkoski data tell us the number of genera *B*_*i*_ that originated in that interval. In our model, we imagine that every genus *µ* comes into the world with a built-in “toughness” *x*_*µ*_. This represents the ensemble of traits that might make a genus more resistant to perturbations, reduced for simplicity to a single number. In our model, for each epoch, we create *B*_*i*_ genera and assign each of them a random toughness number, drawn from some distribution *P* (*x*).

Now consider an extinction event. The Sepkoski data tell us how many genera go extinct at that event. Call this number *D*_*i*_. Our model then asks: what level of environmental *stress S*_*i*_ would be needed to kill exactly *D*_*i*_ genera, given the toughnesses of the genera that are alive at that moment? In the simplest model, we postulate that the stress wipes out the weakest genera first, and we choose the toughness threshold *S*_*i*_ so that precisely *D*_*i*_ are removed. Under this rule, the specific choice of the toughness distribution *P* (*x*) is irrelevant; any choice can be mapped into any other choice by reparameterizing toughness (only rank-order matters). In our simulations, we use a log-normal distribution with unit width in log space.

This framework builds on Newman’s stress-threshold model [14, 15]. However, Newman’s formulation sought to explain the statistics of extinction events, drawing stress threshold magnitudes from a distribution, and treating extinction-count statistics as an output. In contrast, our approach takes the empirical values *B*_*i*_, *D*_*i*_ (total number of originations and extinctions assigned to each geologic interval) as input, and generates a set of *S*_*i*_ (the environmental stresses of the events separating geologic intervals) as well as a synthetic instance of a fossil record (Fig. 2A). By construction, this synthetic dataset will match the empirical *B*_*i*_ and *D*_*i*_ exactly. The other statistics, including the shape of the event survival curves, constitute a prediction of the model.

**FIG. 2.**
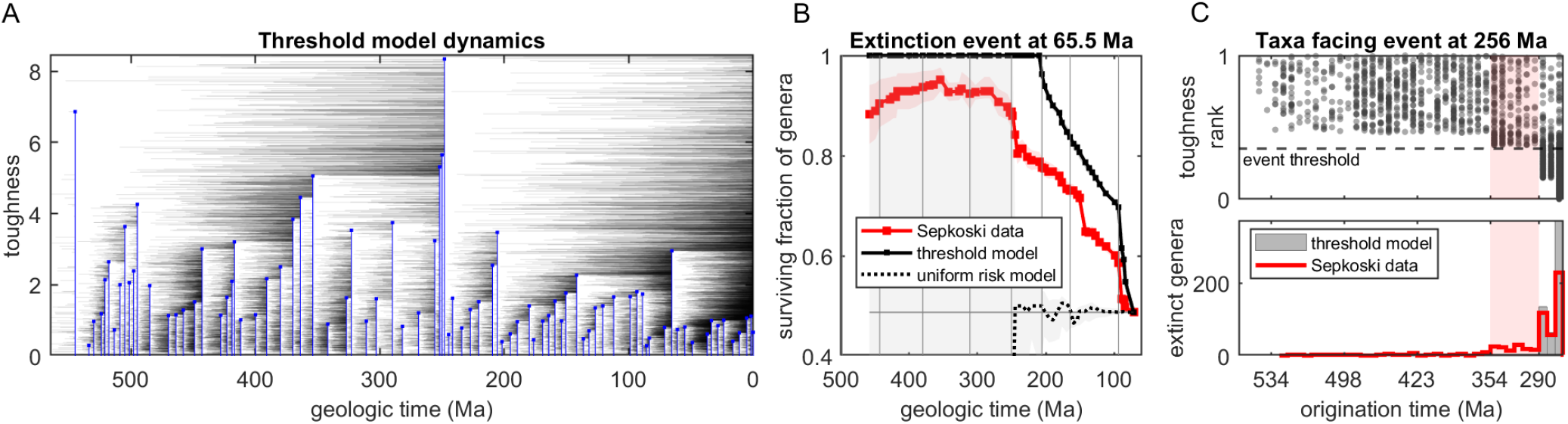
A zero-parameter model reproduces survival advantage of old genera. A: One synthetic fossil record from the toughness-threshold model. Genera are assigned a random “toughness” (gray horizontal lines); epoch boundaries are modeled as stress thresholds (blue lines) chosen so as to match the empirical number of extinctions. Because high-threshold events remove low-toughness genera, the surviving pool becomes enriched in tougher lineages without any explicit age-dependent change in toughness. B: Event survival curve for the K-Pg extinction event at 65.5 Ma. Sepkoski data are shown in red squares; the threshold model is the solid black line. For contrast, the dotted line shows the uniform extinction risk model, in which deaths are sampled uniformly among extant taxa. Shading indicates the IQR computed across the 20 replicates of the models (light gray) or across 1000 bootstrap resamplings of empirical genera (light red). The threshold model captures the qualitative rise in survival probability with genus age (compare to the uniform-risk model, which is flat). However, the deterministic model overstates the survival advantage of old cohorts. C: The mechanism of this failure for one illustrative model realization at the event at 256 Ma. Top, genera alive before the event are plotted by origination time and toughness rank (scaled so that least / most tough map to 0 and 1 respectively); the dashed line is the deterministic death threshold, below which model genera are killed. Bottom, the origination-time distribution of killed genera in the same model realization (gray bars) is compared with extinctions recorded in Sepkoski data (red). Under a deterministic model, all genera in the highlighted cohort (light-red shading) are protected. In contrast, the empirical record does include extinctions from those older cohorts, motivating a stochastic extension of the model (“luck factor”).

An alternative zero-parameter rule could be to select the *D*_*i*_ genera randomly and uniformly among all genera currently alive. This protocol also matches *B*_*i*_ and *D*_*i*_ exactly, but as we will see, the statistics generated by this model will be quite different. This scenario is the law of constant extinction [13] (no notion of toughness), and it will serve as a natural comparison for the toughness-based model.

### C. Survivorship bias is sufficient to reproduce qualitative trends, but overstates the survival advantage of old genera

Fig. 2B shows an example event survival curve (the K-Pg event), overlaid with the predictions of the two models just defined. The rightmost tips of all three curves coincide by construction: this value is the overall survival probability in the extinction event computed across all taxa that experience it, and is fully determined by birth/death statistics. However, the rest of the curve differs.

Under the uniform extinction risk model, the taxa to become extinct are selected uniformly at random among all those alive; as a result, the extinction probability has no age-dependence and the event survival curve is flat (Fig. 2B, dotted black curve), except for sampling stochasticity. (Note that since this model lacks a mechanism to promote the survival of old genera, beyond a certain cohort age the shaded error bars diverge, as the corresponding cohorts become empty. When this happens, the dotted line departs the flat expectation marked with the horizontal gray line.) In contrast, the toughness model captures the age-dependent trend whereby older genera are more likely to survive the focal extinction event (Fig. 2B, solid black curve).

The trends shown for the K-Pg event are representative of the trends observed for other extinction events. Across the ensemble of event survival curves, the model explains 25% of variance, with no free parameters. By construction, this is achieved without any explicit time-dependence of genus properties, but purely through the survivorship bias mechanism.

While the toughness model qualitatively reproduces the age-dependent trend, it systematically overstates the advantage of old taxa. This stems from modeling extinction events as deterministic thresholds, whereby surviving a past extinction *guarantees* survival in future extinctions of lesser magnitude. This is overly simplistic, and comparing model predictions to the data provides strong motivation to relax this assumption. As an illustration, consider Fig. 2C. The top panel shows the scatter plot of toughness versus age across all genera about to experience the extinction event at 256 Ma, for one realization of the model. The Sepkoski dataset instructs us to remove 585 genera, which sets the stress threshold as indicated by the dashed line in Fig. 2C (top). Note that some older taxa (with origination dates up to 354 Ma; highlighted) have toughness barely above the cutoff. Under the deterministic threshold rule, they are protected, and the taxa marked for extinction in the model all originate no earlier than 290 Ma. However, the empirical age distribution of extinct taxa disagrees (Fig. 2C, bottom panel), revealing that a small fraction of older taxa does, in fact, go extinct. This example suggests that a probabilistic survival rule would greatly improve the agreement with the empirical record.

### D. A one-parameter extension predicts age-conditioned survival fractions with high accuracy

The observations of Fig. 2 motivate us to extend our model with one parameter *σ* controlling the extent of stochasticity in the filtering (“luck factor”). At each extinction event, we let every genus draw a temporary multiplicative factor—a bit of good or bad fortune—from another log-normal distribution parameterized by the log-space width *σ*. The effective toughness of this genus at that event is the intrinsic toughness multiplied by this luck factor. Genera are then rank-ordered by the effective toughness, and the stress threshold is selected to precisely match the extinction count as before. When *σ* = 0, there is no luck, and we recover the previous model (Fig. 2B, solid black line). When *σ* is very large, luck dominates; extinction becomes independent of previous history, and we recover the uniform extinction risk model (Fig. 2B, dotted black line). The intermediate values of luck interpolate between these two cases.

Note that unlike the deterministic-threshold model, this model variant is no longer invariant under changes of the shape of toughness distribution. Here we use a log-normal distribution as the simplest choice of a distribution with positive support (see Methods for details).

Figure 3A shows the dynamics of this 1-parameter model. Unlike the deterministic case, it is now possible for a genus to have a lower toughness than the threshold, and yet survive due to luck; Fig. 3A highlights one example of a lucky survival for the Permian-Triassic extinction event (green line crossing the thick blue line). Conversely, a genus can also have higher toughness than the stress threshold, and yet go extinct (red line).

**FIG. 3.**
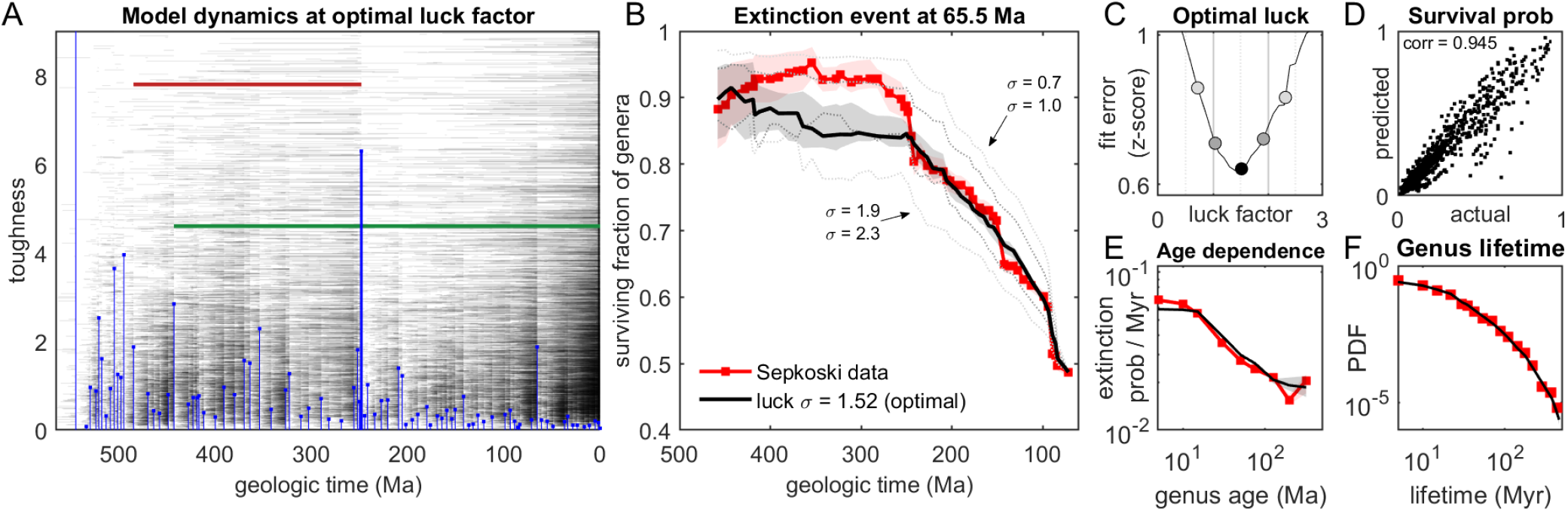
A single parameter makes the survivorship-bias model quantitatively predictive. A: An example synthetic fossil record from the toughness-plus-luck model. Gray segments show genus lifetimes and intrinsic toughness, while blue stems show event thresholds. Unlike in Fig. 2, survival is judged on intrinsic toughness modified by a multiplicative event-specific random luck factor, so genera may survive an event exceeding their intrinsic toughness, or go extinct despite high intrinsic toughness (examples highlighted in green and red, respectively). B: Event survival curve for the same K-Pg event (65.5 Ma) shown in Fig. 2B. Sepkoski data (red) compared to the toughness-plus-luck model prediction at optimal luck factor *σ ≃* 1.52 (solid black) and a few off-optimal luck values (thin dotted curves; see panel C). Shading indicates IQR computed as in Fig. 2B. C: Fit error across the full ensemble of 84 event survival curves as a function of luck factor, measured as the full-curve *z*-RMS discrepancy (see Methods); lower values indicate better agreement. The minimum occurs near *σ* ≃ 1.52. The marked off-optimal points correspond to the dotted curves in (B). D: Predicted versus actual age-conditioned survival fractions across all event-age bins with more than 20 empirical genera exhibit an excellent correlation (*r* = 0.945). Predictions are medians across 20 model replicates. E: Aggregate extinction probability per Myr as a function of genus age. Sepkoski data in red; optimal toughness-plus-luck model in black. Shading indicates IQR as in B; for most points, IQR is too small to be visible. F: Genus lifetime distribution. Colors and shading as in E.

The performance of the toughness-plus-luck model is shown in Fig. 3B-F. Fig. 3B illustrates how the model fits the same event survival curve as previously shown in Fig. 2B. The toughness-plus-luck model curve is shown for 5 values of luck (see Fig. 3C). The middle value (solid black line) uses *σ* = 1.52, which is the optimal fit to the ensemble of the event survival curves (see Methods for details); this is also the value used in panel A. To help interpret this optimal magnitude of the luck factor, we compute the fraction of all extinction events (all pairs (*µ, i*) such that genus *µ* went extinct in event *i*) where the extinction was due to “bad luck” (*x*_*µ*_ *> S*_*i*_; red line in Fig. 3A). For luck factor *σ* = 1.52, about half of extinctions in the model-generated synthetic record are due to bad luck (57.3 ± 0.1%, mean ± SD over 20 replicates).

At this optimal value, the model captures age-conditioned survival probabilities with correlation 0.95 (Fig. 3D; 64.3% of variance explained). As a corollary, the coarser aggregate properties such as the overall dependence of genus extinction probability on age (Fig. 3E), and the distribution of genus lifetimes (Fig. 3F) are captured near-perfectly (for the most part, within error bars of binomial sampling). We conclude that a substantial fraction of the statistical features of the fossil record can be captured by a simple one-parameter model based solely on survivorship bias, with no explicit age dependence of genus properties.

### E. Introducing an explicit time-dependence does not improve fit

We can now return to the biological question posed in the Introduction. There are many reasons to expect the extinction resistance of a genus to change through time. Adaptation, ecological entrenchment, expansion in geographic range, or within-genus diversification could all make older genera systematically harder to eliminate. However, an increase in survival probability with genus age is not, by itself, diagnostic of such within-lineage change: as the models above show, the same pattern can arise from survivorship bias even when each genus has a fixed intrinsic toughness. We are now in a position to ask: do the event survival curves contain evidence for a dataset-wide, directional change in genus properties beyond the survivorship bias already captured by our null model? To test this, we extend the model to include both survivorship bias and a constant rate of toughness growth over time, and ask whether this additional time dependence improves the fit. Specifically, we extend the model by introducing a toughness growth rate *λ* and assume that the toughness of genus *i* at time *t* is 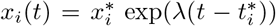, where 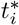 is the start of the epoch in which the Sepkoski dataset records genus *i* to have originated, and 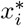 is the initial toughness (drawn randomly as before). The parameter *λ* is a global parameter that applies equally to all taxa, and we will allow both positive and negative values, corresponding to a consistent toughness increase or decline over time, respectively. This exponential dependence is the simplest choice consistent with positivity of *x*_*i*_ – though as we will see, the relevant values of *λ* will be very small, so an exponential and a linear model would behave similarly.

This model has an extra fitting parameter (for a total of two), and an extra degree of freedom should of course be able to provide a better fit. However, the quality of the optimal fit is improved only marginally (Fig. 4), and crucially, the optimal value of *λ* is remarkably small (|*λ*_opt_| < 0.001; see Sup. Fig. S3). To interpret the scale of the y axis in Fig. 4, it is convenient to think about toughness in log space. With our modeling choices (a multiplicative luck factor *σ* distributed log-normally), survival in a given extinction event is determined by a stochastic draw whose random component (in log space) is drawn from a normal distribution of width *σ* ≃ 1.5. For a median-length epoch of *T*_median_ = 5.75 Ma, this stochastic contribution is to be compared with *λ T*_median_ (the deterministic trend accumulated over the same period), setting the relevant scale of *λ* to *σ/T*_median_ = 0.26 units of log-toughness per Ma. Thus, the entire range of the y axis in Fig. 4 corresponds to deterministic effects that are at least an order of magnitude weaker than the stochastic contribution.

**FIG. 4.**
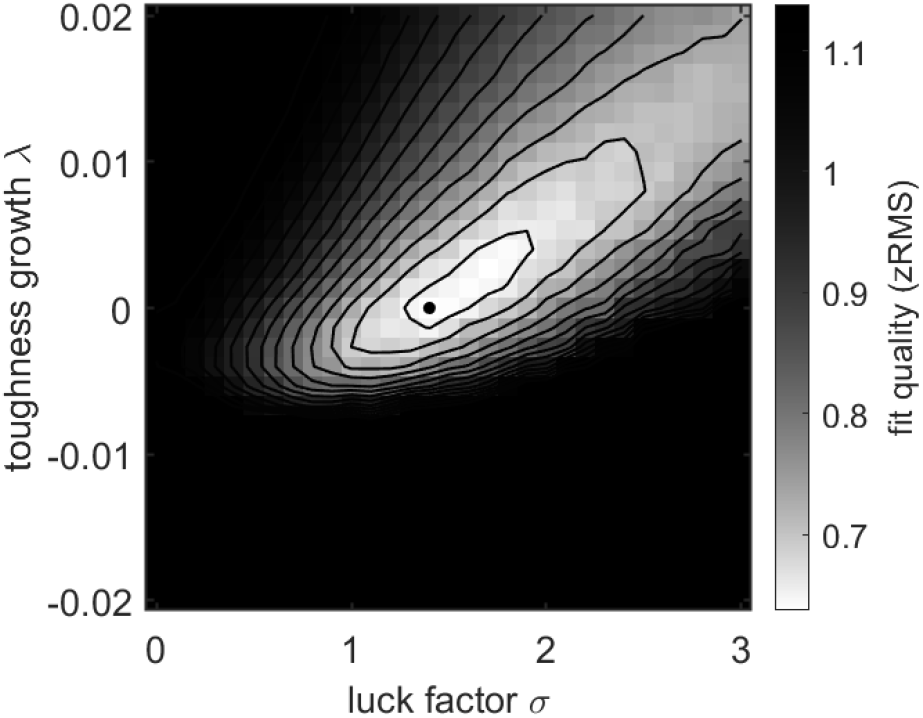
Allowing toughness to consistently increase or decrease over time does not improve the fit. The heat map shows the full-curve *z*-RMS fit error for a two-parameter extension where we vary both the luck factor and the rate of systematic toughness growth (positive or negative). Each grid point is averaged over 10 synthetic model replicates; lighter shading indicates lower error, and black contours mark equal-error levels. The heat map shows a coarse parameter grid. The optimum (black dot) lies very close to zero growth (|*λ*_opt_| < 0.001; see Sup. Fig. S3), indicating that a consistent increase or decrease of intrinsic toughness through genus lifetime is not needed to explain the event survival curves.

To summarize, even when time-dependence is allowed to be a free fitting parameter, the best fit to the data is a model in which the time-dependent trend is negligible compared to the stochastic effect of the luck factor. We conclude that at least from a statistical perspective (in the aggregate across all genera), an explicit time-dependent entrenchment does not appear to be a significant effect.

### F. The role of a null model is to highlight outliers

The previous sections characterized the overall fit quality of the model. In this section, we ask whether this fit quality is uniform across the dataset, or whether the analysis identifies some specific events as outliers worthy of attention.

For any given event, the model sets up an expectation for which genera are more likely to be eliminated, based on their prior survival history. For each genus alive before the event, we can infer a posterior distribution over toughness, and estimate its probability of going extinct in a synthetic extinction event constrained to kill the same number of genera as the empirical event. We can then define an event weirdness score, which assesses whether the set of eliminated taxa is unusually surprising given this fixed-extinction-count comparison (see Methods). The constraint on empirical extinction count ensures that our weirdness score is not directly sensitive to event magnitude: whether an event eliminates many genera or few, we only ask if their *identities* are unusual (see Sup. Fig. S5).

The results are shown in Fig. 5. We see that most empirical events are no more unusual than synthetic events generated from the model itself (dark gray band). However, event indices 31, 37, 41, and 43 stand out as unusually positive. These correspond to significant genus turnover pulses at the Tournaisian–Viséan boundary (∼342 Ma), the Asselian–Sakmarian boundary (∼282 Ma), the Dzhulfian–Dorashamian interval (∼250 Ma), and the Induan–Olenekian boundary (∼245 Ma), with the ∼250 Ma pulse forming part of the prolonged end-Permian extinction crisis.

**FIG. 5.**
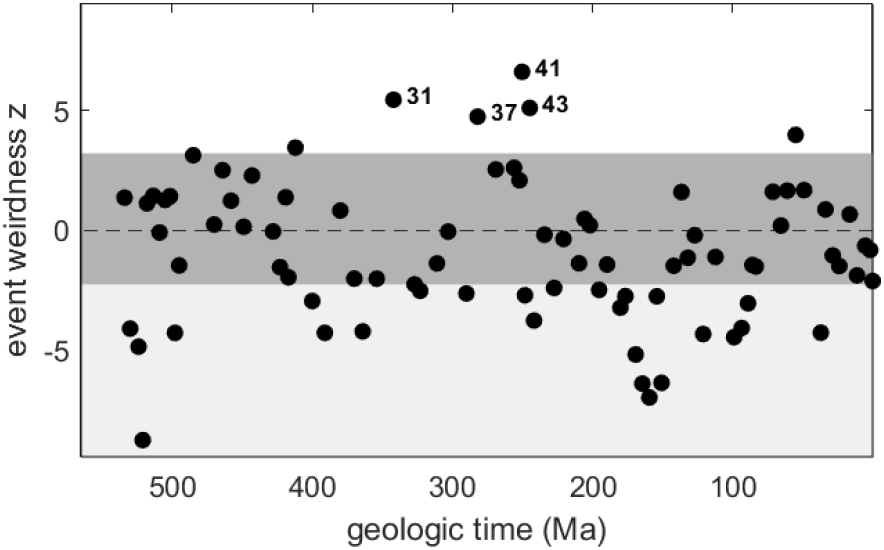
The fitted null model isolates a small number of unusually structured extinction events. Black points indicate the weirdness *z*-score for each empirical extinction event. The weirdness score quantifies whether the empirical cohort of extinct taxa appears unusual under the toughness-plus-luck model, given the Bayesian posterior on toughness inferred for each genus from its prior survival history; see methods for details. For reference, the dark gray band is the middle 99% of weirdness score range computed across 20 synthetic fossil records generated by the fitted toughness-plus-luck model (which by definition is compatible with itself). Positive values lying above this reference band indicate events that eliminated an unusual set of taxa; the negative-z outliers (light gray) are discussed in the text. Most empirical events fall within the expected range, but event indices 31, 37, 41, and 43 stand out as unusually positive.

Fig. 5 also shows many events with large negative *z*-scores (light gray band). These deserve a comment. Large negative *z*-scores indicate cases where the extinct cohort is in unusually strong agreement with the toughness ranking. To understand how this is possible, recall that the large fitted luck factor corresponds to strong stochasticity. Thus, an event where the extinct taxa are too strongly enriched in low-toughness representatives will also score as unusual. One way to interpret this is that fitting a single luck factor across all 84 events likely overestimates the stochasticity of some events, which is not necessarily surprising. The strong positive outliers are of much greater interest, as these are the cases where the lists of extinct genera actually *contradict* the toughness order inferred by the model, and might be biologically informative, indicating events that “play by different rules”. For instance, the Asselian–Sakmarian boundary and the Induan–Olenekian boundary (events 37 and 43) are both relatively mild (with only 12% and 14% extinction fraction, respectively), yet eliminate an unusually large number of very old genera. This directly contradicts the model expectations, suggesting that the stress applied in these events probed a different notion of genus resilience than most other extinction events, and highlighting them as natural targets for more detailed biological analyses.

## DISCUSSION

In this work, we investigated how much of the observed age dependence of extinction can be explained by survivorship bias alone. We find that a simple toughness-plus-luck model with a single fitted parameter reproduces the ensemble of age-conditioned survival probabilities remarkably well, along with the age dependence of extinction rates and the genus lifetime distribution. Introducing an explicit time-dependent increase in genus toughness provides no additional explanatory power. We therefore find no evidence that a systematic, dataset-wide trend in genus toughness needs to be postulated to explain the overall age dependence of extinction risk. We emphasize, however, that this conclusion concerns the aggregate structure only: our analysis does not exclude age-dependent changes within individual lineages that leave no imprint on the pooled statistics. Distinguishing these would require lineage-resolved comparisons.

It is worth being clear about what we mean by “luck.” At the most direct level, luck represents genuine contingency: even very similar genera need not share the same fate in a given extinction event. However, luck also plays a second, more subtle role. Our model deliberately compresses all biological determinants of extinction susceptibility into a single latent toughness. It is clear that in reality, glaciation, volcanism, ocean acidification, bolide impacts, and other perturbations probe different combinations of physiology, ecology, life history, and geographic range. The fitted luck factor therefore captures not only intrinsic stochasticity, but also the inevitable mismatch between this multidimensional biology and its one-dimensional summary.

This compression defines the model’s scope: by ignoring taxonomic identity, ecology, physiology, abundance, geographic distribution, and environmental context, we do not aim to describe the fossil record completely. Rather, we ask how much of its statistical structure can be captured while invoking none of these ingredients. The answer is that one parameter accounts for roughly 64% of the variance in age-conditioned survival probabilities. The events that deviate most strongly from this baseline (Fig. 5) are those that eliminate genera hardest to reconcile with a single toughness axis. Our hope is that this framework gives paleontologists an additional quantitative tool for identifying the features of the fossil record that are especially likely to be biologically informative.

## METHODS

### A. Sepkoski data loading and preprocessing

We analyzed genus-level stratigraphic ranges from a late electronic version of J. J. Sepkoski Jr.’s marine fossil genus database, provided by Sepkoski to one of the authors and used with his permission. This database forms the basis of the subsequently published Compendium of Fossil Marine Animal Genera [7], which we cite as the archival reference for the dataset.

For each genus, the Sepkoski dataset assigns its origination and extinction time to a stratigraphic interval. For our analysis, we therefore reason in terms of epoch boundaries (the “extinction events” separating stratigraphic intervals). For simplicity, below, we use the shorthand of describing originations and extinctions as occurring ‘at’ these boundaries. Specifically, for an event *E* separating intervals *A* and *B*, we will refer to the genera originating in *B* as the “births” assigned to *E*, and the genera going extinct during *A* as the “deaths” assigned to this event. Thus, “originated at t” always means “the Sepkoski database assigns the origination to the stratigraphic interval directly following t”, but this shorthand simplifies presentation.

To assign numerical ages to the stratigraphic interval labels, we considered using the publicly available R package `sepkoski` [16], which provides an updated numerical timescale. However, inspection of that package revealed a number of apparent inconsistencies in the interval-to-age conversion, and we therefore chose to perform this conversion ourselves directly from the original database using the International Stratigraphic Chart 2000 [17]. We selected this contemporaneous timescale to maximize consistency with the stratigraphic terminology used in Sepkoski’s database and to minimize reinterpretation of the original interval assignments according to subsequent revisions of stratigraphic nomenclature. A derived, human-readable version of the processed data, together with all analysis scripts needed to reproduce every figure, is included in the public archive accompanying this work.

A small amount of additional preprocessing was required because some adjacent boundaries in the source table split a single biological or stratigraphic transition into two nearly coincident numerical entries. In several such cases, one boundary records births but no deaths, while the other records deaths but no births. Although this convention is harmless for many analyses, it is problematic here because our model treats each boundary as a distinct extinction opportunity. Thus, if births are assigned to the earlier of two adjacent boundaries and deaths to the later, genera originating during the transition are treated as having been exposed to—and survived—an additional extinction event. Thus, splitting a single transition into separate birth and death boundaries introduces artifacts.

To remove these artifacts, prior to computing all survival curves and model fits, we eliminated all epoch boundaries recorded as corresponding to zero births or zero deaths, by merging them with the closest adjacent boundary. (The first and last epoch boundaries are an exception, and were not merged despite having zero deaths and zero births, respectively.) For each merge pair (*a, b*), all appearances and disappearances assigned to boundary *a* were reassigned to boundary *b*. As a consistency check, prior to merging, we confirmed that the dataset contained no genera recorded as originating at one member of the pair and disappearing at the other.

### B. Choice of toughness and luck distributions

In the deterministic threshold model, the choice of toughness distribution is immaterial because only the rank ordering of genera enters the dynamics. Any continuous toughness distribution can therefore be transformed into any other by a monotone reparameterization, leaving all model predictions unchanged. We use a log-normal distribution simply as a convenient representative.

This invariance no longer holds once stochastic luck is introduced, because survival depends on the numerical values of toughness rather than their ordering alone. We therefore model both intrinsic toughness and the event-specific luck factor as log-normal random variables. Besides being the simplest choice with positive support, this makes the multiplicative survival rule additive in logarithmic coordinates, where both intrinsic toughness and luck are represented by Gaussian variables. This representation substantially simplifies the posterior inference of genus toughness from survival history used in the event-weirdness analysis (Fig. 5).

### C. Model fitting and optimization

All model fits were performed against the full ensemble of event survival curves. For each focal event *E* and an earlier boundary *t* (filtering event), the empirical curve entry 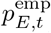 is the fraction of genera that survive event *E*, among genera that were alive at *t* and lived to see *E* (i.e., were alive both at *t* and immediately before *E*). For a candidate model and parameter value, we generated multiple synthetic fossil records constrained to have the same per-boundary numbers of births and deaths as the empirical data. For each synthetic record, we computed the same ensemble of event survival curves. Model predictions were summarized by the median synthetic value across replicates, and residuals were defined as

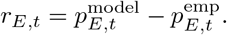

Fitting raw probability residuals is not ideal because different points on the survival curves are estimated with very different precision. Points based on small retrospective cohorts have large sampling error and should not be given the same weight as points based on hundreds of genera. We therefore fit in standardized residual space. For each empirical curve entry with cohort size *N*_*E,t*_, we estimated the binomial sampling uncertainty using a Jeffreys-regularized survival probability,

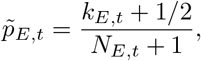

where *k*_*E,t*_ is the number of genera in the cohort that survived the focal event. The corresponding binomial standard error is

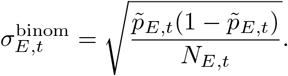

At the same time, a pure binomial-error weighting would over-trust the very largest cohorts. The model is not intended to capture every taxonomic, environmental, sampling, and stratigraphic detail of the data, so once an empirical survival probability is already estimated well enough, increasing the cohort size further should not give that point arbitrarily large leverage. We account for this by adding a precision-crossover scale, *ϵ*, to the denominator:

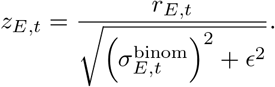

In the analyses shown here, *ϵ* = 0.05. This parameter can be interpreted as the residual scale below which other sources of model-data mismatch dominate over binomial sampling error. Equivalently, it specifies a “good enough” cohort size. When 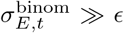, the point is low precision and is downweighted by its sampling error. When 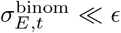, the point is already precise enough and receives approximately the same weight as other high-precision points. For survival probabilities near one half, *ϵ* = 0.05 corresponds to a crossover cohort size of approximately 1*/*(4*ϵ*^2^) ≃ 100 genera.

The scalar fit objective was the root-mean-square of these standardized residuals over all included points on all event survival curves,

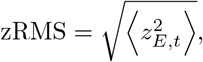

where the average is taken over finite curve entries passing the cohort-size and event-validity masks. Note that since the cohorts defining a single event survival curve are nested and therefore not statistically independent, this quantity (zRMS) is a discrepancy metric rather than a formal likelihood. Its purpose is to compare model variants using the same weighted summary of their residual structure.

The one-parameter toughness-plus-luck model was fitted by numerically minimizing this full-curve zRMS over the luck factor *σ*; accordingly, the y axis in Fig. 3 reports the quality of fit in zRMS units. For reference, in raw probability units, the RMS deviation between the optimal-luck model and the empirical event survival curves is 0.06 (i.e., survival probabilities are typically predicted within 6 percentage points), and can be visually assessed using Fig. 3D.

For the two-parameter model in Fig. 4, we first evaluated the full-curve *z*-RMS on a coarse grid of luck factor and toughness-growth rate, which is the grid shown in the heat map. To place the optimum more precisely, we then also evaluated a refined 31 × 31 grid centered on the minimum of the coarse grid, with tenfold finer spacing in each parameter (see SI and the Sup. Fig. S3). We used grid refinement rather than a continuous optimizer because, for a fixed set of stochastic model replicates, the simulated extinction sets change only when variation in the parameters alters the rank ordering of effective toughness among taxa. The resulting objective is therefore piecewise constant, and a grid search is more robust. The contour lines in Fig. 4 are guides for the eye (computed from a Gaussian-smoothed version of the coarse-grid heat map), and were not used in the analysis in any way.

### D. Event weirdness score

The event weirdness analysis asks whether the identities of the genera eliminated in a given empirical extinction event are unusual under the assumptions of the toughness-plus-luck model, conditional on the observed number of extinctions in that event. This conditioning ensures the score is not a measure of extinction intensity, but of whether the identities of the eliminated genera appear surprising given the incoming cohort and the number of extinctions.

We work in log-toughness units. Let *q*_*u*_ denote the median stress threshold estimated for event *u* (across the fitted model replicates). For a genus of intrinsic toughness *x*, the probability of going extinct in event *u* is

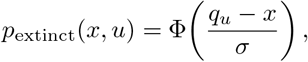

where Φ is the standard normal cumulative distribution function, and *σ* is the luck factor, which we set to *σ* = 1.52 as per our fitting.

For each focal event *t*, we first inferred a posterior distribution over toughness for every genus in the incoming cohort, using only its survival history prior to the focal event. Specifically, for a genus first observed in epoch *f* and alive immediately before event *t*, the posterior is proportional to

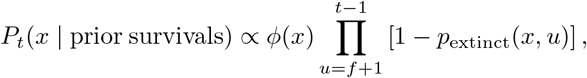

where *ϕ*(*x*) is the standard normal prior density. Thus, genera that have survived earlier high-threshold events acquire posterior distributions shifted toward higher toughness. The focal event itself is not included in this posterior; its extinctions are what the weirdness score evaluates.

Let *C*_*t*_ be the incoming cohort for event *t*, and let *D*_*t*_ be the empirical number of genera in this cohort that disappear at that event. To construct the fixed-*D*_*t*_ null distribution, we performed 1000 Monte Carlo draws. In each draw, we sampled a toughness value for every genus in *C*_*t*_ from its posterior distribution, added an independent luck term of width *σ*, and recorded the *D*_*t*_ genera with the lowest resulting bids. This produces a Monte Carlo ensemble of extinction sets, all with the same total number of extinctions as the empirical event.

From these Monte Carlo extinction sets, we estimated the model-predicted extinction probability 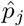for every genus *j* ∈ *C*_*t*_, equal to the fraction of Monte Carlo extinction sets in which that genus was included. To avoid infinite scores from finite Monte Carlo sampling, probabilities were clipped to the interval [1*/*(2*M*), 1 − 1*/*(2*M*)], where *M* = 1000 is the number of Monte Carlo draws. Each genus was then assigned an extinction-surprise score

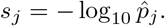

The observed event weirdness statistic was defined as the total surprise of the empirically extinct genera,

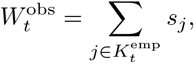

where 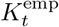is the empirical extinction set. We compared this value to the corresponding statistic computed for each Monte Carlo extinction set,

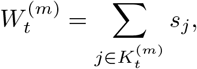

and reported the standardized weirdness score

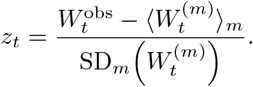

Positive values therefore indicate events whose empirical extinction set contains genera that the fitted model expected to be unusually likely to survive. Negative values indicate the opposite: extinction sets that are highly concentrated among model-vulnerable genera, more so than expected under the fitted value of luck.

As a calibration check, we applied the same event-weirdness procedure to synthetic fossil records generated by the fitted toughness-plus-luck model. For these synthetic records, each replicate’s own event thresholds were used. As expected, when evaluated on synthetic records generated by the model itself, no event looks consistently special and the weirdness profiles are statistically flat; see Fig. S4. The gray reference band in Fig. 5 shows the central 99% interval, computed by pooling finite weirdness *z*-scores across the 20 fitted-model synthetic replicates and all events.

## DATA AVAILABILITY STATEMENT

All analysis was performed in MATLAB (Mathworks, Inc.). Analysis scripts and data files needed to reproduce all figures in this manuscript are publicly available at Mendeley Data [18]. The original historical electronic database provided by Sepkoski cannot be redistributed by the authors because they do not have authorization to do so. However, a derived human-readable data file sufficient to reproduce all the figures in this work is included with the code.

## ACKNOWLEDGEMENTS

Research reported in this publication was supported by the National Institute of General Medical Sciences of the National Institutes of Health under Award Number R35GM160222, and by grants from the NSF (DMS-2235451) and Simons Foundation (MP-TMPS-00005320) to the NSF-Simons National Institute for Theory and Mathematics in Biology (NITMB). The content is solely the responsibility of the authors and does not necessarily represent the official views of the National Institutes of Health. This work was completed at a Kavli Institute for Theoretical Physics (KITP) program, supported by grant NSF PHY-2309135 and the Gordon and Betty Moore Foundation Grant No. 2919.02 to KITP.

## SUPPLEMENTARY INFORMATION

## Appendix A

### Direct comparisons of narrowly defined sister cohorts are statistically noisy

The most direct test of survivorship bias would compare genera originating immediately before and immediately after an earlier filtering event *A*, and ask how well those two cohorts survive a later focal event *B*. In this approach, each cohort is restricted to genera originating in a single interval adjacent to *A* and still alive immediately before *B*. However, as we will see, the small size of these cohorts limits the utility of this approach.

To see this, let’s use these cohorts to test a much simpler hypothesis: that older genera survive better than younger genera. This broad age dependence is well established and clearly visible in the aggregate event survival curves of the main text. If the direct sister-cohort comparisons struggle to recover even this strong effect, they are unlikely to have sufficient power to detect the considerably subtler signal sought in this work.

Fig. S1 applies the direct sister-cohort comparison to every ordered pair of extinction events. For each pair, it compares the survival probabilities of the two cohorts (survive till *B* and originating just before *A*, versus survive till *B* and originating just after *A*). Cell color indicates the posterior evidence for the sign of

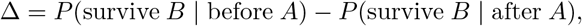

with red supporting greater survival of the older cohort, blue supporting the reverse, pale colors indicating weak evidence, and gray indicating that one or both conditioned cohorts are empty.

Overall, the matrix is reassuringly biased toward red, consistent with the well-established tendency of older genera to survive better. However, many comparisons are undefined, very few individual comparisons provide strong evidence, and those that do are concentrated near the diagonal, where the two cohorts have experienced relatively little attrition before the focal event. Thus, even the broad age-dependence visible in the aggregate event survival curves is only weakly recovered by direct sister-cohort comparisons. Establishing the much subtler signal sought in this work—that filtering events produce survival differences beyond those expected from ordinary age dependence—would therefore be even more challenging. This motivates the nested retrospective cohorts used in the main text.

## Appendix B

### Discontinuities in event survival curves are not explained by cohort age alone

Event survival curves in Fig. 1 show pronounced discontinuities near some earlier extinction events. One possible explanation is purely compositional: a major extinction may sharply alter which genera remain in the retrospective cohort, producing a corresponding change in the cohort’s mean age.

Supplementary Fig. S2 evaluates this possibility by plotting the mean genus age at the focal event for the same retrospective cohorts used to construct the three example survival curves in Fig. 1. The mean-age curves change much more smoothly than the corresponding event survival curves, including across major earlier extinction events. Thus, the discontinuities in Fig. 1 cannot be explained solely by changes in cohort age.

## Appendix C

### Refined parameter search around the optimum

Figure S3 shows a local refinement of the parameter search around the minimum identified by the coarse grid in Fig. 4. At this finer resolution, the objective function exhibits substantial point-to-point variation arising from the finite number of stochastic model replicates.

Although the refined search does not isolate a unique best-fitting point within the low-error basin, it shows that all near-optimal solutions lie very close to zero toughness growth (|*λ*_opt_| ≲ 0.001). As explained in the main text, a deterministic toughness change comparable in magnitude to the event-specific stochastic variation represented by the luck factor would require *λ* ≈ 0.26. Thus, Figure S3 shows that any deterministic change in intrinsic toughness over a genus lifetime is negligible compared to the stochastic variation represented by the luck factor.

## Appendix D

### Calibration of the event-weirdness statistic

Before interpreting empirical event-weirdness scores, it is important to verify that the statistic behaves as expected under the fitted null model. Supplementary Fig. S4 applies the complete event-weirdness calculation (see Methods and main text Fig. 5) to synthetic fossil records generated by the fitted toughness-plus-luck model. The median synthetic weirdness remains close to zero, with no extinction event receiving a consistently elevated or depressed weirdness score. This demonstrates that the event-weirdness statistic is well calibrated under the fitted null model.

## Appendix E

### Independence of event weirdness from extinction magnitude

The event weirdness score introduced in Fig. 5 is intended to quantify *which* genera are eliminated during an extinction event, conditional on the observed number of extinctions. It is therefore designed to assess the composition of the extinction list rather than the overall severity of the event. To verify this property, Supplementary Fig. S5 plots event weirdness against extinction fraction for both the empirical Sepkoski record and synthetic fossil records generated by the fitted toughness-plus-luck model. As expected, the correlation between event weirdness and extinction fraction is weak. Panel A highlights the four events flagged in the main text, confirming that the unusually structured events flagged by this analysis are not simply the largest extinction events.

**FIG. S1.**
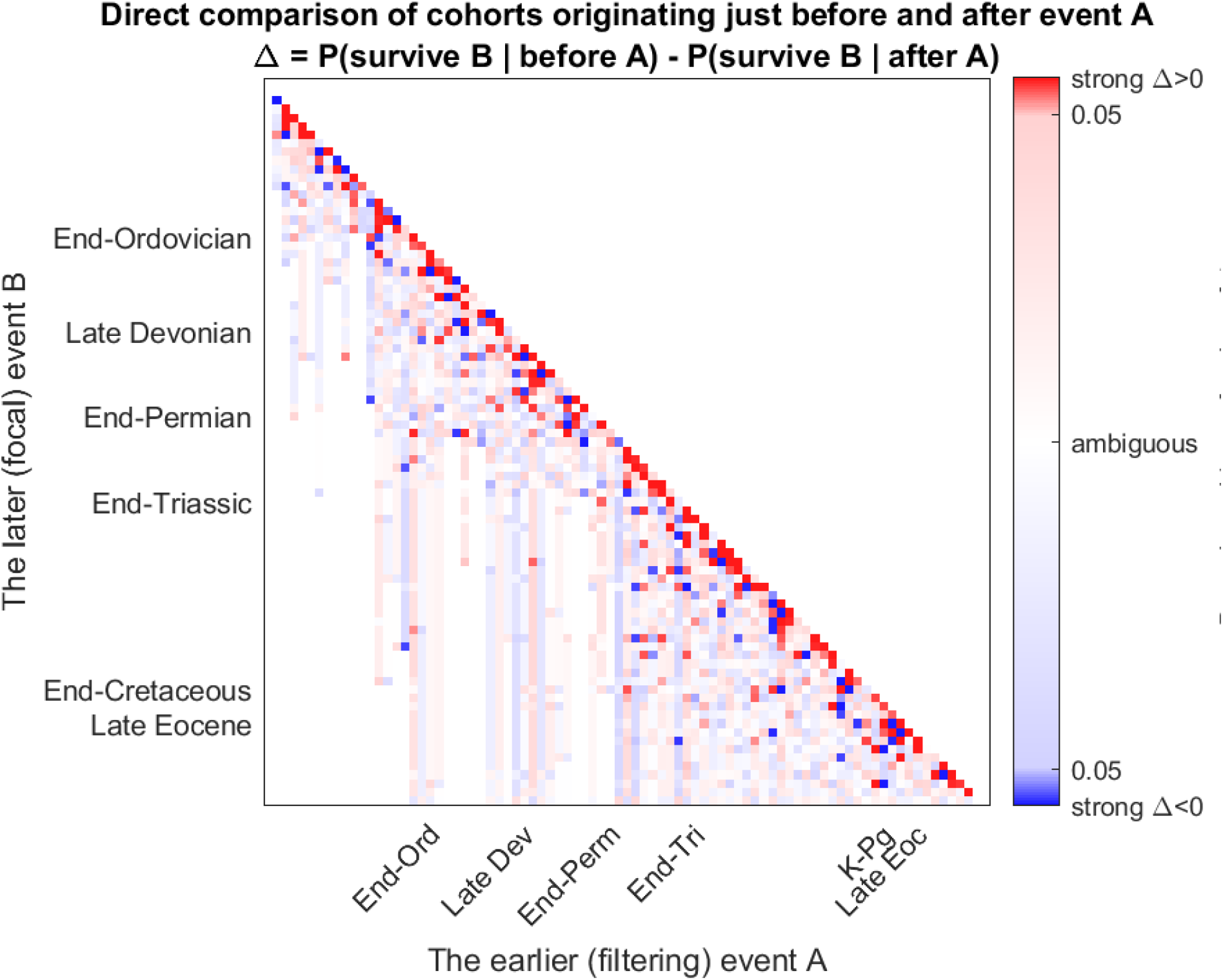
Direct sister-cohort comparisons provide limited statistical power. Each pixel compares the survival probability through a later (focal) extinction event *B* of genera originating immediately before and immediately after an earlier (filtering) event *A*, conditioned on being alive immediately before *B*. Events *A* and *B* run through all 84 epoch boundaries recorded in the dataset; the ticks on the axes mark the Big Five extinction events and the Late Eocene event. Cell color indicates the posterior evidence for the sign of Δ, where Δ *>* 0 corresponds to greater survival of the older (pre-*A*) cohort. Gray cells indicate comparisons for which one or both conditioned cohorts are empty. Although the matrix shows an overall bias toward positive Δ, the evidence is generally weak and becomes sparse away from the diagonal because of progressive cohort attrition.

**FIG. S2.**
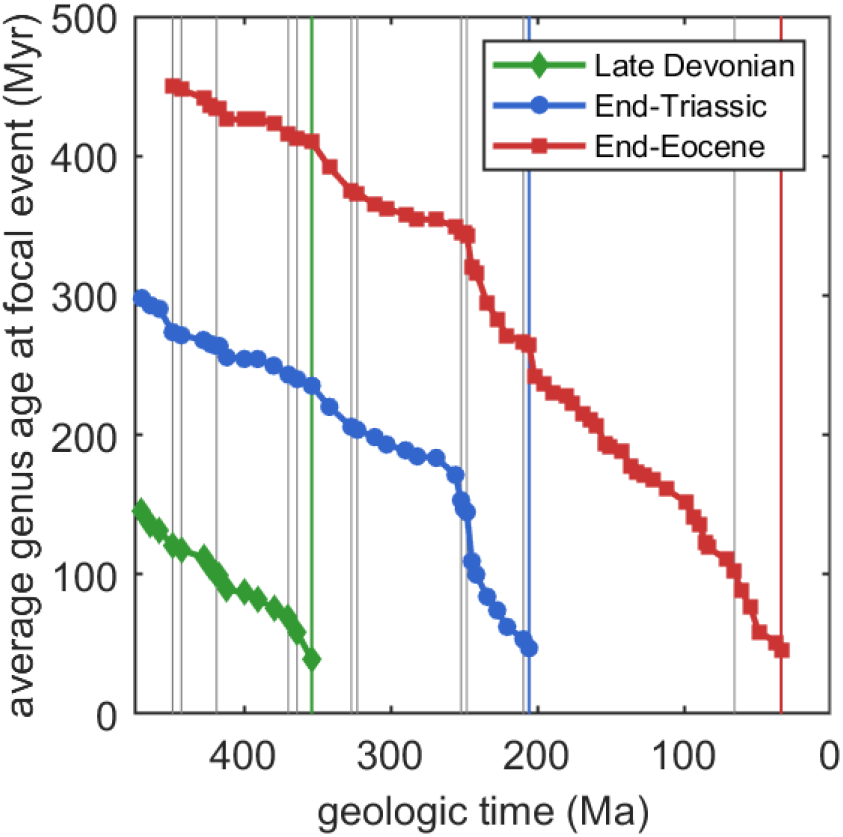
Mean cohort age changes smoothly across earlier extinction events. For the same three focal extinction events shown in Fig. 1C, each curve gives the mean genus age at the focal extinction event among genera contributing to the corresponding point on the event survival curve. Figure layout is identical to Fig. 1C to facilitate visual comparison: gray vertical lines mark earlier extinction events with extinction fractions exceeding 40%, and colored vertical lines mark the focal events. Although major extinction events alter the composition of the retrospective cohorts, the resulting changes in mean cohort age are comparatively smooth. Thus, the sharp discontinuities observed in the event survival curves of Fig. 1C cannot be explained solely by changes in cohort age.

**FIG. S3.**
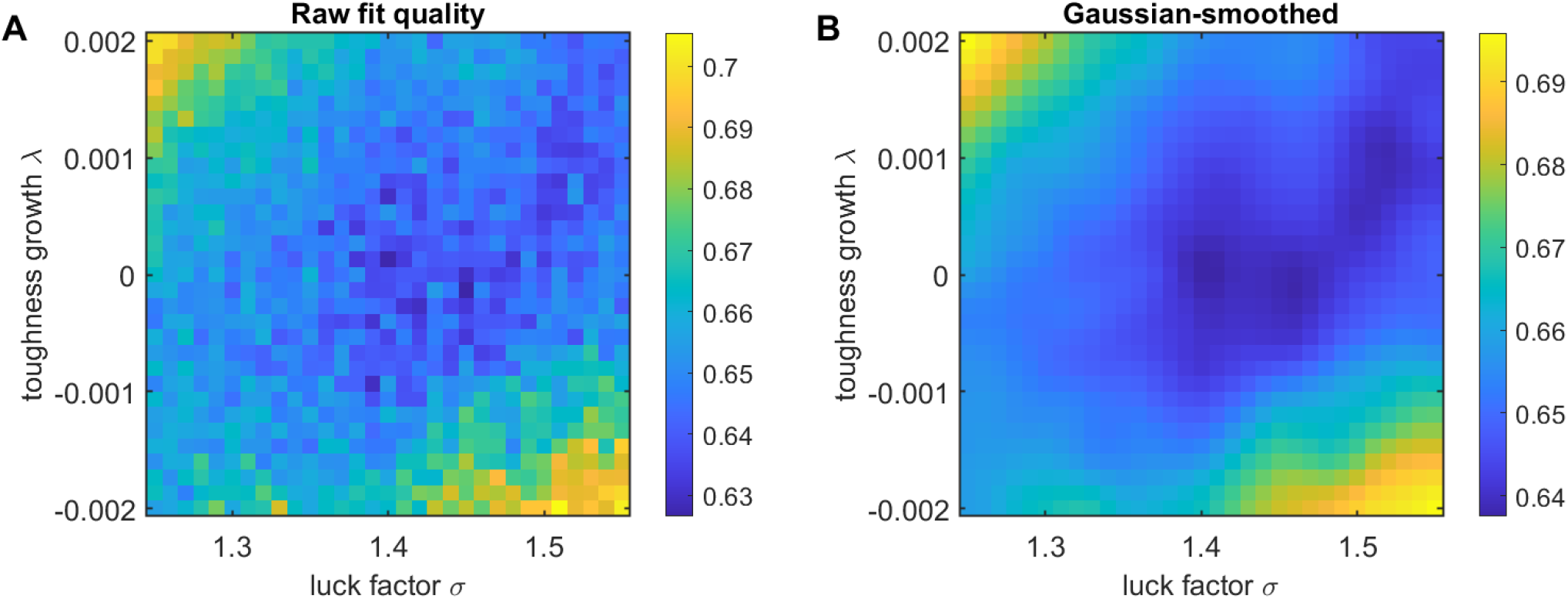
Local refinement of the luck-factor and toughness-growth fit. **A:** Full-curve *z*-RMS fit error evaluated on a 31 × 31 refined parameter grid centered on the minimum of the coarse grid shown in Fig. 4. Each grid point is averaged over the same 10 synthetic model replicates used in the coarse sweep. **B:** The same refined-grid fit surface after modest Gaussian smoothing, shown to highlight the underlying low-error region.

**FIG. S4.**
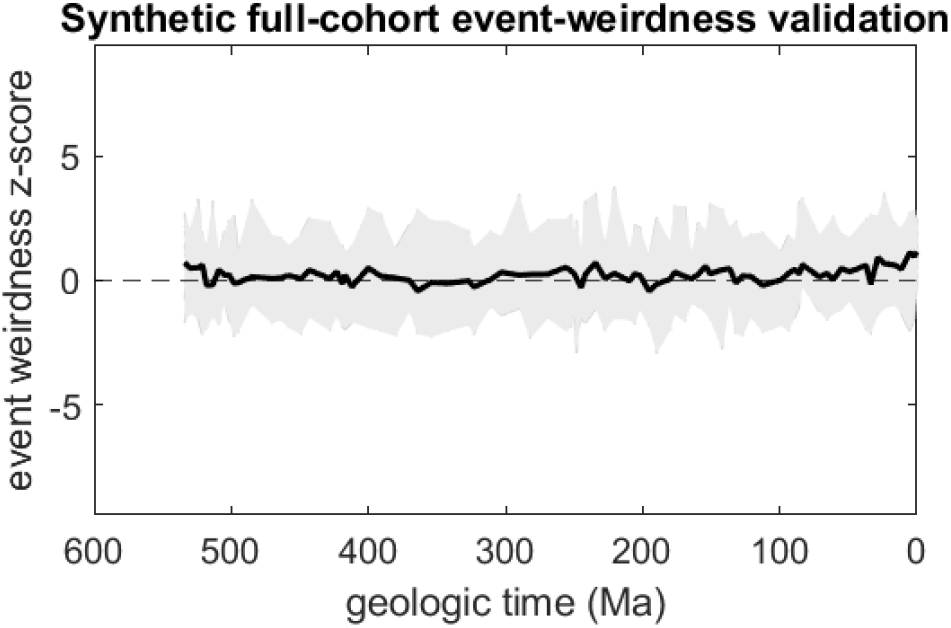
Synthetic validation of the event-weirdness statistic. Event weirdness scores computed from synthetic fossil records generated by the fitted toughness-plus-luck model. The black curve shows the median, and gray band the range of the weirdness score computed across 20 synthetic replicates. The y-axis scale is identical to that of Fig. 5 in the main text. As expected under the null model, synthetic event weirdness fluctuates around zero, with no extinction event consistently exhibiting unusually large positive or negative scores.

**FIG. S5.**
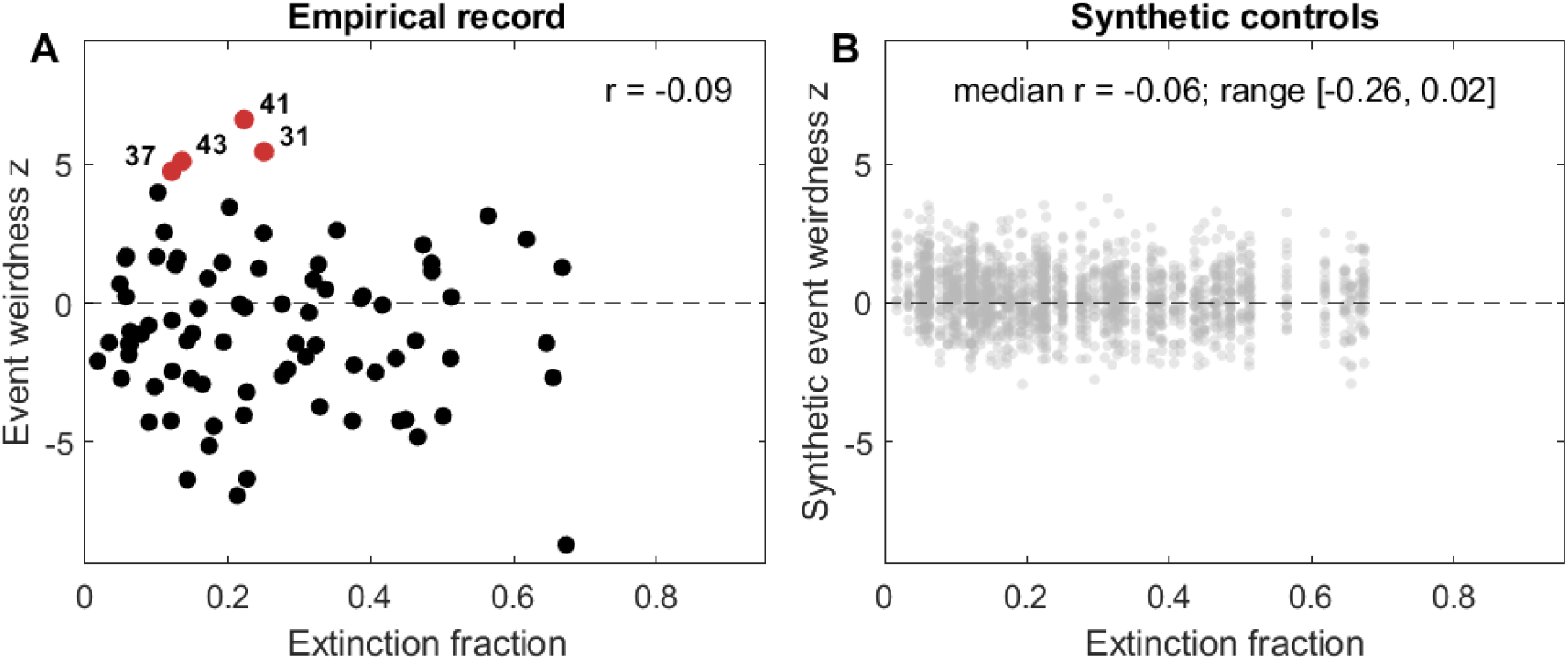
Event weirdness is independent of extinction magnitude. (A) Event weirdness score versus extinction fraction for the empirical Sepkoski record. The four unusually positive events highlighted in Fig. 5 (event indices 31, 37, 41, and 43) are shown in red. The weak correlation (Pearson *r* = *−*0.09) indicates that event weirdness is not simply a consequence of extinction magnitude. (B) The corresponding analysis for synthetic fossil records generated by the fitted toughness-plus-luck model. Each gray point is one event from one synthetic replicate; all replicates share the empirical extinction fractions by construction. Correlations remain weak, confirming that the event weirdness score is effectively decoupled from event magnitude.

